# A non-spike nucleocapsid R204P mutation in SARS-CoV-2 Omicron XEC enhances inflammation and pathogenicity

**DOI:** 10.1101/2025.05.28.656516

**Authors:** Shuhei Tsujino, Masumi Tsuda, Jumpei Ito, Sayaka Deguchi, Taha Y. Taha, Hesham Nasser, Lei Wang, Julia Rosecrans, Rigel Suzuki, Saori Suzuki, Kumiko Yoshimatsu, Melanie Ott, The Genotype to Phenotype Japan (G2P-Japan) Consortium, Terumasa Ikeda, Kazuo Takayama, Kei Sato, Shinya Tanaka, Tomokazu Tamura, Takasuke Fukuhara

**Affiliations:** Department of Virology, Faculty of Medicine Sciences, Kyushu University, Fukuoka, Japan; Department of Microbiology and Immunology, Faculty of Medicine, Hokkaido University, Sapporo, Japan; Department of Cancer Pathology, Faculty of Medicine, Hokkaido University, Sapporo, Japan; Institute for Chemical Reaction Design and Discovery (WPI-ICReDD), Hokkaido University, Sapporo, Japan; Division of Systems Virology, Department of Microbiology and Immunology, The Institute of Medical Science, The University of Tokyo, Tokyo, Japan; Center for iPS Cell Research and Application (CiRA), Kyoto University, Kyoto, Japan; Department of Synthetic Human Body System, Medical Research Institute, Institute of Integrated Research, Institute of Science Tokyo; Gladstone Institutes, San Francisco, California, United States of America; Bioengineering and Therapeutic Sciences, University of California, San Francisco, California, United States of America; Division of Molecular Virology and Genetics, Joint Research Center for Human Retrovirus Infection, Kumamoto University, Kumamoto, Japan; Institute for Vaccine Research and Development (IVReD), Hokkaido University, Sapporo, Japan; Institute for Genetic Medicine, Hokkaido University, Sapporo, Japan; Department of Medicine, University of California, San Francisco, California, United States of America; Chan Zuckerberg Biohub – San Francisco, San Francisco, California, United States of America; AMED-CREST, Japan Agency for Medical Research and Development (AMED), Tokyo, Japan; Graduate School of Frontier Sciences, The University of Tokyo, Chiba, Japan; Graduate School of Medicine, The University of Tokyo, Tokyo, Japan; International Research Center for Infectious Diseases, The Institute of Medical Science, The University of Tokyo, Tokyo, Japan; International Vaccine Design Center, The Institute of Medical Science, The University of Tokyo, Tokyo, Japan; Collaboration Unit for Infection, Joint Research Center for Human Retrovirus infection, Kumamoto University, Kumamoto, Japan; MRC-University of Glasgow Centre for Virus Research, Glasgow, UK; One Health Research Center, Hokkaido University, Sapporo, Japan; Laboratory of Virus Control, Research Institute for Microbial Diseases, The University of Osaka, Suita, Japan

**Keywords:** SARS-CoV-2, COVID-19, XEC, nucleocapsid, R204P, pathogenicity, inflammation, NF-κB

## Abstract

The global circulation of SARS-CoV-2 in human populations has driven the emergence of Omicron subvariants, which have become highly diversified through recombination. In late 2024, SARS-CoV-2 Omicron XEC variant emerged from the recombination of two JN.1 progeny, KS.1.1 and KP.3.3, and became predominant worldwide. Here, we investigated virological features of the XEC variant. Epidemic dynamics modeling suggested that spike substitutions in XEC mainly contribute to its increased viral fitness. Additionally, four licensed antivirals were effective against XEC. Although the fusogenicity of XEC spike is comparable to that of the JN.1 spike, the intrinsic pathogenicity of XEC in hamsters was significantly higher than that of JN.1. Notably, we found that the nucleocapsid R204P mutation of XEC enhanced inflammation through NF-κB activation. Recent studies suggest that the evolutionary potential of spike protein is reaching its limit. Indeed, our findings highlight the critical role of non-spike mutations in the future evolution of SARS-CoV-2.

## INTRODUCTION

RNA viruses are prone to high mutation rates, so environmental changes can lead to the emergence of progeny viruses that are resistant to drugs and immune defenses. Genetic variation in RNA viruses typically occur through substitution, deletion and insertion. In addition, dramatic variation through recombination occurs in RNA viruses and is thought to have played a significant role in the recent evolutionary histories.^1–4^ A notable example is severe acute respiratory syndrome coronavirus 2 (SARS-CoV-2),^5^ the causative agent of the highly contagious disease COVID-19. Because humans experience infection and/or vaccination, the circulating variants, namely Omicron have evolved to exhibit reduced intrinsic pathogenicity, increased transmissibility, and enhanced immune escape compared with ancestral variants.^6–15^ The continuous circulation of Omicron has led to the emergence of "recombinant variants" through simultaneous infection with multiple variants and recombination in the host. Omicron XBB lineage was the most prevalent lineage worldwide, a recombinant between BA.2 subvariants BJ.1 and BM.1.1.1 (a descendant of BA.2.75). XBB is the first SARS-CoV-2 variant whose fitness increased through recombination rather than substitution.^11^ Other recombinant variants, such as XBC.1.6 (Delta variant B.1.617.2 × BA.2) and XDD.1.1 (EG.5.1.1 × JN.1), have also been identified, though they did not become the predominant.

Omicron XEC variant emerged from the recombination of two JN.1 descendants, KS.1.1 (JN.13.1.1.1) and KP.3.3 (JN.1.11.1.3.3). XEC was first identified in Germany in August 2024, has rapidly spread in several Western contenents.^16^ As of April 2025, XEC are predominantly circulating worldwide according to Nextstrain (clade 24F; https://nextstrain.org/ncov/gisaid/global/6m). Because XEC is a chimera of KS.1.1 and KP.3.3 variants, spike (S) protein of XEC inherited key mutations enhanced binding affinity to the angiotensin-converting enzyme 2 (ACE2) receptor from KP.3.3 and acquired mutations for the immune-evasive properties from KS.1.1.^16–20^ This suggests that XEC S maintains a similar affinity to ACE2 and a higher capacity to evade immunity compared with the previously dominant variants, KP.3 and KP.3.1.1. In addition, XEC has the R204P mutation in the nucleocapsid (N) protein and the A419T mutation in the NSP2 protein, which it inherited from KP.3.3 and KS.1.1, respectively. In the past variants, the co-occurring mutations at positions 203 and 204 of N (R203K/G204R) is involved in the efficient assembly of viral particle, leading to increased viral replication and pathogenicity of SARS-CoV-2.^21–23^ The effect of the N:R204P mutation on the virological characteristics including replication and pathogenicity has not been reported.

In this study, we aimed to characterize the virological features of SARS-CoV- 2 XEC variant and investigate how the R204P mutation in the nucleocapsid protein affects viral replication and pathogenicity.

## RESULTS AND DISCUSSION

### Mutations contributing to the increased viral fitness of XEC

Compared with JN.1, XEC has five amino acid substitutions (S:T22N, S:F59S, S:F456L, S:Q493E and S:V1104L) in S and two substitutions in the non-spike protein genes (NSP2:A419T and N:R204P).^16^ To identify the mutations contributing to the rapid spread of XEC, we applied a hierarchal Bayesian multinomial logistic model established in our previous study to viral genome surveillance data from GISAID (https://www.gisaid.org/).^10^ Here, fitness refers to the relative effective reproductive number (Re) between variants, estimated under the assumption that the relative values of the Re among variants remain constant over time. This model predicts the effect of mutations by assuming that the fitness of a given variant is represented as the sum of the effects of its constituent mutations.^10^ When multiple mutations are highly co-occurring, it becomes challenging to estimate their individual effects. To address this, these co-occurring mutations were grouped into mutation clusters, and their effects were estimated at the mutation cluster level. We applied this model to viral genome epidemiological surveillance data from the USA, covering the period from January 1, 2024, to December 31, 2024. This dataset includes 209 mutation clusters, composed of 277 distinct mutations **(Table S1)**.

Among the 209 analyzed mutation clusters, 15 were estimated to contribute positively to increased fitness (**Figures 1A and 1B**). The mutation cluster that exhibited the highest positive effect corresponded to mutations acquired on the stem branch of the BA.2.86 lineage or the JN.1 lineage. Notably, among the seven characteristic mutations of XEC, S:F456L, S:Q493E, S:F59S, and S:T22N were estimated to have a significant positive effect on fitness, ranking in the top 2, 3, 5 and top 8, respectively. In contrast, no statistically significant effect was detected for S:V1104L, NSP2:A419T and N:R(G)204P (**Figure 1A**). These results suggest that the rapid spread of XEC was driven by S:F456L, S:Q493E, S:T22N, and S:F59S rather than other substitutions.

**Figure 1.**
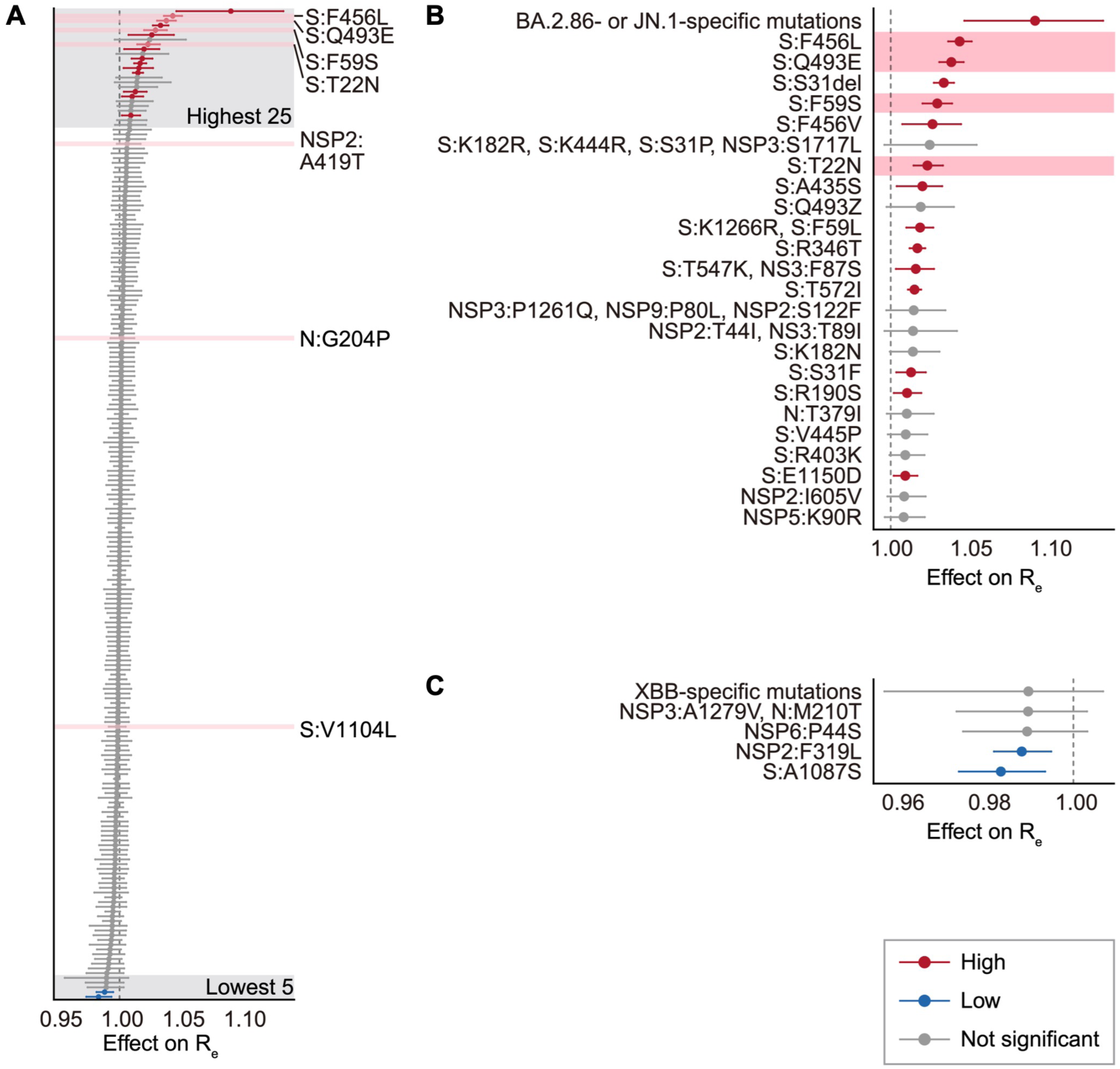
Estimation of effects of mutations on viral fitness **(A)** Estimated effect of each mutation on relative Re estimated by a hierarchal Bayesian model. The posterior mean value (dot) and 95% credible interval (line) are shown. A group of highly co-occurred mutations (e.g., those acquired on the stem branch of the BA.2.86 or JN.1 lineage) was treated as mutation clusters. The red and blue dots indicate the substitutions with significant positive and negative effects, respectively. Mutations characteristic to XEC are highlighted in pink. **(B-C)** Estimated effect of each mutation on relative Re. Panel **(B)** shows the top 25 mutations with the highest estimated effects, while panel **(C)** presents the 5 mutations with the lowest estimated effects.

### Fusogenicity of XEC S

The fusogenicity of the XEC S protein was measured by the SARS-CoV-2 S protein-mediated membrane fusion assay using Calu-3/DSP1-7 cells (**Figure S1A**).^24^ The surface expression level of XEC S was comparable to that of the parental JN.1 S (**Figure S1B**). As previously reported,^24–26^ the Delta S protein exhibited the greatest fusogenicity, while the KP.3 S protein exhibited the weakest fusogenicity (**Figure 2A**). In case of XEC S, comparable fusogenicity to the JN.1 S protein was observed, suggesting the XEC S protein contributes similar viral pathogenicity to the JN.1 S protein.

**Figure 2.**
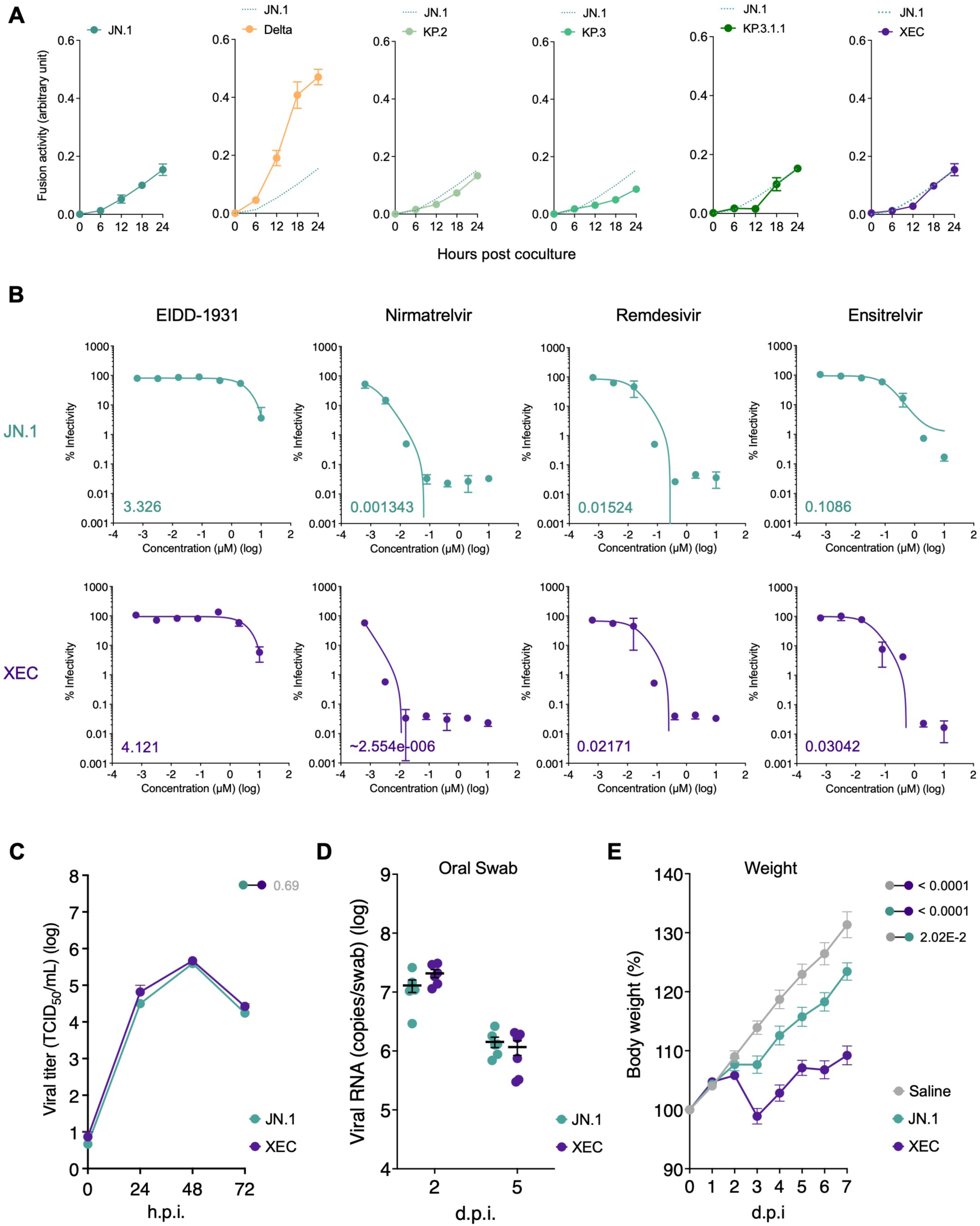
Virological features of the SARS-CoV-2 XEC **(A)** S-based fusion assay in Calu-3 cells. The dashed green line indicates the result of JN.1. The red number in each panel indicates the fold difference between JN.1 and the derivative tested at 24 h post coculture. Assays were performed in quadruplicate. Statistically significant differences versus JN.1 across time points were determined by multiple regression. **(B)** Effect of antiviral drugs against XEC. Antiviral effects of the four drugs (EIDD-1931, nirmatrelvir [also known as PF- 07321332], remdesivir, and ensitrelvir) in human iPSC-derived lung organoids. The assay of each antiviral drugs was performed in triplicate, and the 50% effective concentration (EC50) was calculated. **(C)** JN.1 and XEC were inoculated into VeroE6/TMPRSS2 cells (MOI = 0.01). The 50% tissue culture infectious dose (TCID50) of the culture supernatant were routinely quantified. **(D** and **E)** Syrian hamsters were intranasally inoculated with JN.1 and XEC. Six hamsters of the same age were intranasally inoculated with saline (uninfected). (**D**) Six hamsters per group were quantified viral RNA load in oral swab by RT-qPCR. (**E**) Six hamsters per group were used to routinely measure the body weight. Uninfected (saline) hamster data is also shown. The familywise error rates (FWERs) calculated using the Holm method are indicated in the figures. h.p.i: hours post-infection; d.p.i: days post-infection.

### Antiviral effect of clinically available compounds against XEC

We assessed the sensitivity of XEC to four licensed antiviral drugs, EIDD- 1931, nirmatrelvir (also known as PF-07321332), remdesivir, and ensitrelvir. Clinical isolate of JN.1 was used as a control. Both viruses were inoculated into human induced pluripotent stem cell (iPSC)-derived lung organoids, a physiologically relevant model, and treated with the four antiviral drugs. Among them, nirmatrelvir showed the strongest antiviral effects, with no significant differences in antiviral efficacy were observed between JN.1 and XEC (**Figure 2B**). Remdesivir and ensitrelvir demonstrated significant antiviral effects on the two isolates, whereas EIDD-1931 exhibited moderate antiviral effects on the two isolates.

### Viral replication and pathogenicity of XEC variant

To investigate the replication efficiency and intrinsic pathogenicity of XEC, we used the clinical isolates of JN.1 and XEC. First, we inoculated clinical isolates of JN.1 and XEC into VeroE6 cells expressing TMPRSS2^27^ and quantified viral infectious titers in supernatants. In VeroE6/TMPRSS2 cells (**Figure 2C**), the replication efficiency of XEC was comparable to that of JN.1. Next, clinical isolates of JN.1 and XEC were respectively intranasally inoculated into hamsters, the established animal model for evaluation^6–15,25^. To evaluate viral spread in infected hamsters, we routinely measured the viral RNA load in oral swabs. The viral RNA load of XEC-infected hamsters was comparable to that of JN.1-infected hamsters at 2 and 5 d.p.i. (**Figure 2D**). Interestingly, the body weights of the hamsters infected with XEC were significantly lower than those of JN.1-infected hamsters (**Figure 2E**), suggesting that intrinsic pathogenicity of XEC are higher than that of JN.1.

### Mutation dynamics of nucleocapsid protein

Because N protein functions as the basis for viral RNA genome packaging into ribonucleotide complex (RNP) and assembly into virus particles, functional changes lead to alter viral pathogenicity *in vivo*.^22,28–30^ To investigate the effects of mutation in N, the frequency of mutations in JN.1 and XEC was examined. P13L, E31del, R32del, S33del, G204R, and S413R mutations were acquired in a convergent manner during the evolution of the Omicron variant (**Figure 3A**).^31^ In addition, Q229K mutation was acquired during evolution to BA.2.86.^14^ The previous study showed R203K/G204R mutation contributed to enhance both viral replication and pathogenicity. Therefore, the R204P mutation in XEC may affect alteration of viral replication and pathogenicity.

**Figure 3.**
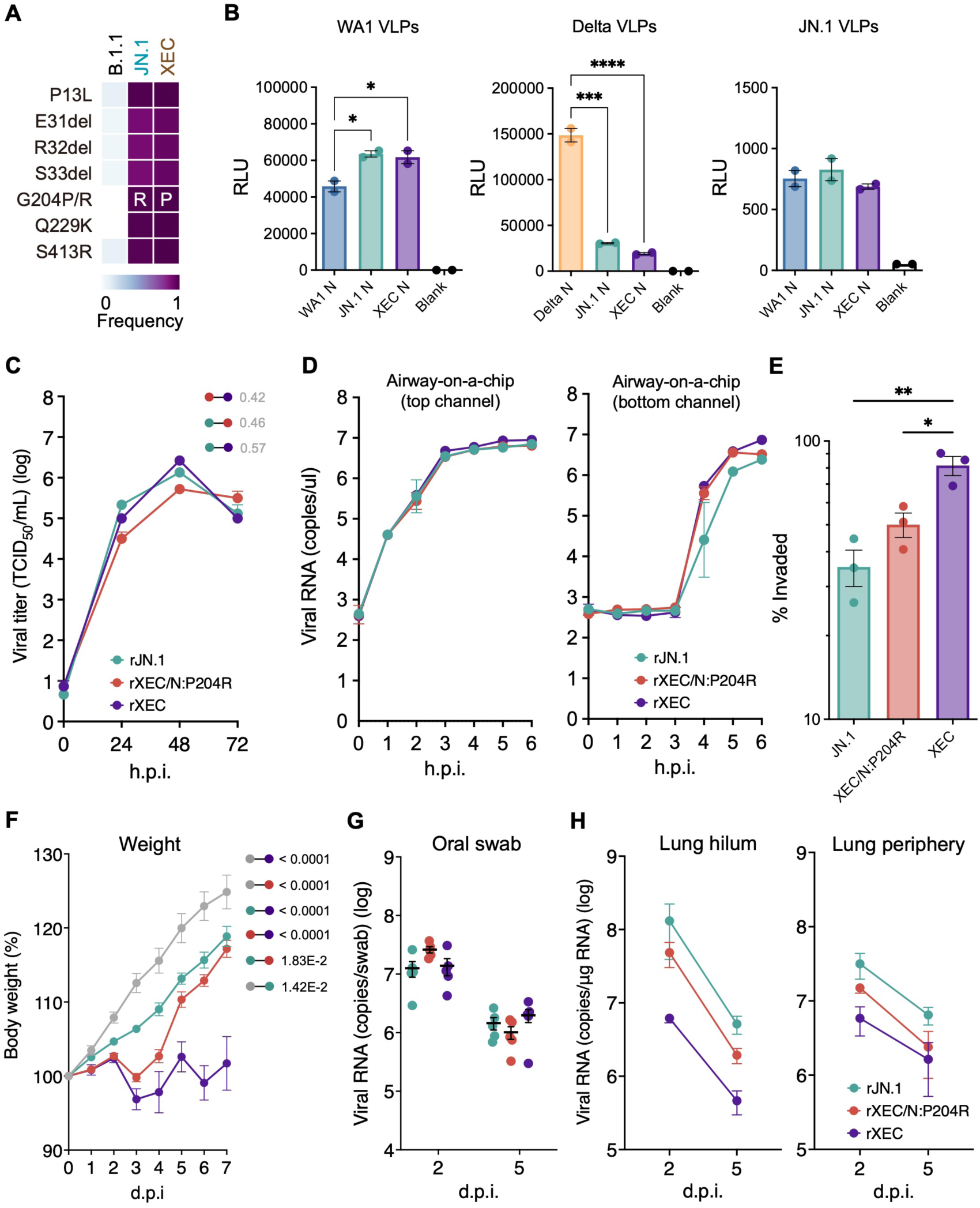
The impact of nucleocapsid R204P mutation on replication and pathogenicity **(A)** Frequency of N protein’s mutations in XEC and other lineages of interest. Only mutations with a frequency >0.5 in at least one representative lineage are shown. **(B)** The efficiency of VLP assembly was measured for different N- protein species indicated and quantified in relative luminescence units. **(C)** Growth kinetics of rJN.1, rXEC/N:P204R, and rXEC were inoculated into VeroE6/TMPRSS2 cells (MOI = 0.01) (left) and Calu-3 cells (MOI = 0.1) (right). The 50% tissue culture infectious dose (TCID50) of the culture supernatant were routinely quantified. **(D)** Recombinant viruses were inoculated into an airway- on-a-chip system. The copy numbers of viral RNA in the top and bottom channels of an airway-on-a-chip were routinely quantified by RT-qPCR (left). The percentage of viral RNA load in the bottom channel per top channel at 6 d.p.i. (i.e., % invaded virus from the top channel to the bottom channel) is shown (right). **(E)** The percentage of viral RNA load in the bottom channel per top channel at 6 d.p.i. (i.e., % invaded virus from the top channel to the bottom channel) is shown. **(F-H)** Syrian hamsters were intranasally inoculated with the recombinant viruses. Six hamsters per group were used to routinely measure the respective parameters. **(F)** Body weight of infected hamsters (n = 6 per infection group). Uninfected hamster data is also shown. **(G)** Viral RNA loads in the oral swab (*n* = 6 per infection group) at 2 and 5 d.p.i. (**H**) Viral RNA loads in the lung hilum (left) and lung periphery (right) of infected hamsters (n = 4 per infection group) at 2 and 5 d.p.i. The FWERs calculated using the Holm method are indicated in the figures.

### Effects of N:R204P on the viral assembly and replication

To evaluate the effect of the N:R204P mutation on the viral assembly, we performed a virus-like particle (VLP) assay. The formation of VLPs in 293T cells containing SARS-CoV-2 structural proteins (S, E, M, and N) and packaging RNA is detected through expression of a luciferase reporter in infected receiver cells.^32,33^ The N:R204P mutation slightly reduced the assembly of VLPs on the Delta backbone, but did not affect that of VLPs on the WA1 or JN.1 backbone (**Figure 3B**). Next, to examine the effect of N:R204P mutation on viral replication, we generated recombinant viruses carrying the single mutation in the N protein of XEC, rXEC/N:P204R, rXEC, and rJN.1. The recombinant viruses were inoculated into the cell cultures we examined as above. In VeroE6/TMPRSS2 cells (**Figure 3C**), no significant differences in growth kinetics were observed between the viruses. These findings suggest that the N:R204P mutation has little effect on the viral replication.

To investigate the impact of XEC infection on the airway epithelial and endothelial barriers, we employed an airway-on-a-chip system. Virus penetration from the upper channel to the lower channel serves as an indicator of the viruses’ ability to breach these barriers. The proportion of viruses that penetrated the lower channel of the XEC-infected airway-on-a-chip was higher compared with JN.1- and XEC/N:P204R-infected airway-on-a-chip (**Figures 3D and 3E**). SARS-CoV-2 N activates human endothelial cells through Toll-like receptor 2 (TLR2)/NF-κB and mitogen-activated protein kinase (MAPK) signaling pathways.^34^ Therefore, N:R204P mutation may affect the airway epithelial and endothelial barriers through endothelial activation.

### Effects of N:R204P on the viral pathogenicity

To further investigate the intrinsic pathogenicity of the three recombinant viruses, the viruses were respectively inoculated as above. Consistent with the result of clinical isolates, the body weights of the hamsters infected with rXEC were significantly lower than those of rJN.1-infected hamsters (**Figure 3F**). Notably, the weight of hamsters infected with rXEC was significantly lower than those of hamsters infected with rXEC/N:P204R, indicating that N:R204P mutation contributes to enhanced pathogenicity.

To evaluate viral spread of the three recombinant viruses in hamsters, the viruses were respectively inoculated into hamsters. We routinely measured the viral RNA load in oral swabs. The viral RNA load of rJN.1-, rXEC/N:P204R- and rXEC-infected hamsters were comparable (**Figure 3G**). We then compared viral spread in respiratory tissues. We collected the lungs of infected hamsters at 2 and 5 d.p.i., and the harvested tissues were separated into the hilum and periphery regions. The viral RNA loads of rXEC-infected hamsters was significantly lower than those of rJN.1- and rXEC/N:P204R-infected hamsters (**Figure 3H left and right**).

### Effects of XEC N:R204P on immunopathogenic features of XEC

We further performed IHC analysis targeting the viral N protein in the respiratory tissues of infected hamsters. At 2 d.p.i, N-positive cells were more detected in the bronchi/bronchioles of rJN.1-infected hamsters than rXEC- and the N:P204R-infected hamsters (**Figure 4A**).

**Figure 4.**
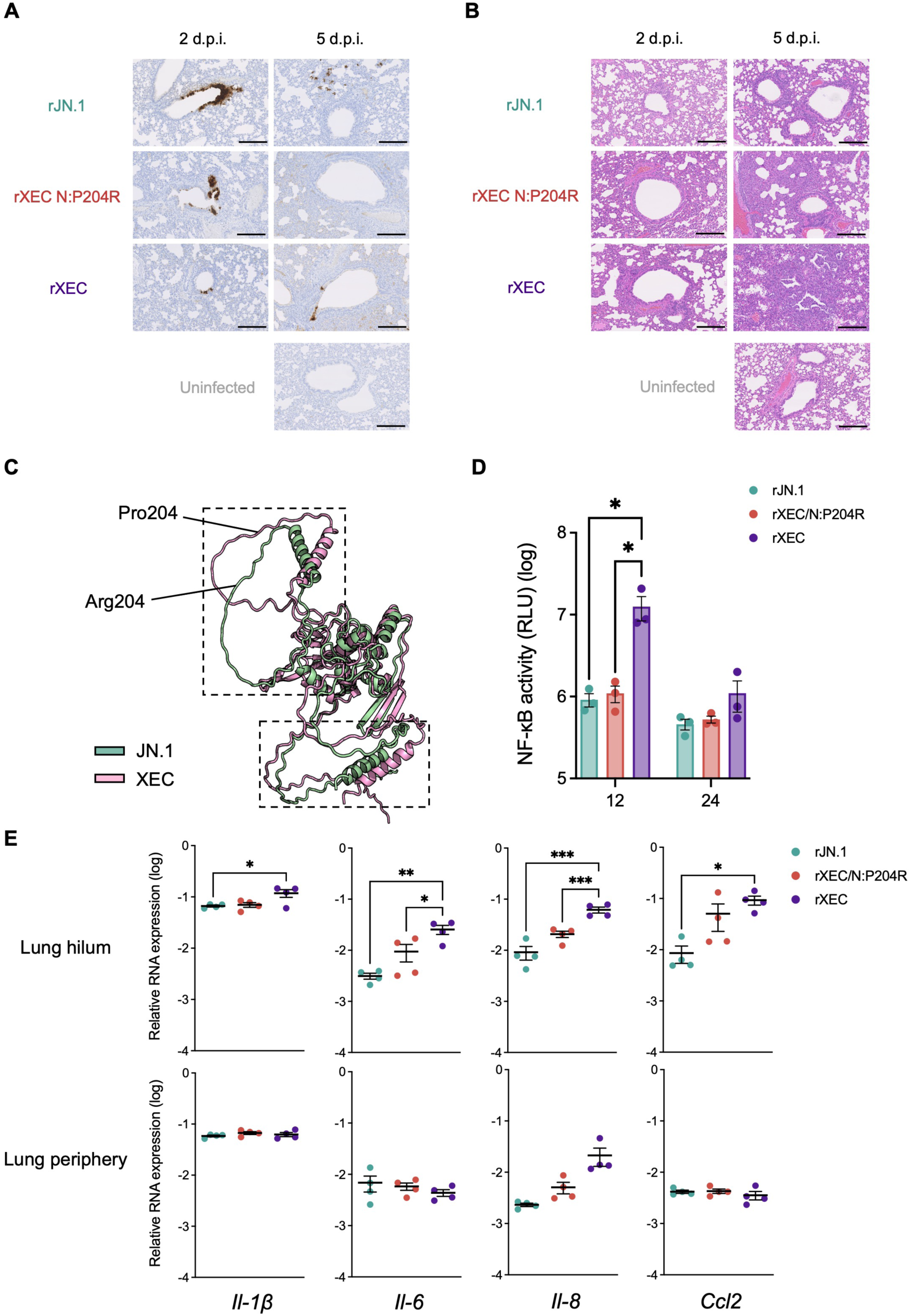
Nucleocapsid R204P mutation enhanced inflammation by NF-κB activation **(A)** IHC of the viral N protein in the lungs at 2 d.p.i. (left) and 5 d.p.i. (right) of infected hamsters. Representative figures (N-positive cells are shown in brown). Uninfected hamster data is also shown. **(B)** H&E staining of the lungs at 2 d.p.i. (left) and 5 d.p.i. (right) of infected hamsters. Representative figures and uninfected lung alveolar space are shown. (**C**) The structure of N (JN.1 N and XEC N) was predicted by Alphafold3. **(D)** HEK293/ACE2/TMPRSS2 cells were transfected with NF-κB reporter vector. At 24 h after transfection, cells were infected with rJN.1, rXEC/N:P204R, and rXEC for 1 h. Luciferase activity was measured at 12 and 24 h.p.i. **(E)** mRNA of the lung tissues obtained at 2 d.p.i. was used to measure expression levels of inflammatory genes (*Il-1β*, *Il-6*, *Il-8*, and *Ccl2*) with normalization using the housekeeping gene *Rpl18*.

To investigate the intrinsic pathogenicity of recombinant viruses, histopathological analyses were performed. At 2 d.p.i, alveolar damage around the bronchi was prominent in rXEC- and the N:P204R-infected hamsters (**Figure 4B**). On the other hand, inflammation was limited in bronchi/bronchioles in the hamsters infected with rJN.1 (**Figure 4B**). At 5 d.p.i, the alveolar architecture was more destroyed by alveolar damage and the expansion of type II pneumocytes in rXEC-infected hamsters (**Figure 4B**). The low viral load in XEC-infected lungs may be due to viral clearance caused by an enhanced immune response, judging from the degree of inflammation (**Figure 4A and 4B**). Notably, a strong inflammation in the acute phase was observed in the lungs of rXEC-infected hamsters even at 5 d.p.i. These results suggest that the N:R204P mutation contributes to increased inflammation in the lung.

### NF-κB activation of XEC N protein

SARS-CoV-2 accessory and non-structural proteins, including ORF3a, ORF7a, NSP5, NSP6 and NSP14, have been shown to activate the NF-κB pathway and induce downstream inflammatory cytokines and chemokines.^35–40^ On the other hand, SARS-CoV-2 N protein has been reported to inhibit NF-κB activation and downstream signaling by inhibiting the formation of the TAK1- TAB2/3 complex.^41^ In addition, N protein lacking interaction with the TAK1- TAB2/3 complex has been shown to induce strong inflammation. Since proline in the linker increases structural rigidity,^42^ we hypothesized that the R204P mutation might alter the overall structure of N protein and impair this regulatory function.

Using AlphaFold3^43^, we found that the amino acid substitutions from arginine to proline at position 204 in XEC N altered the position of the α-helix and the linker (**Figure 4C**). As this α-helix has been reported to be important for interaction with the TAK1-TAB2/3 complex, we investigated the effect of N:R204P mutation on NF-κB activation by NF-κB promoter-driven luciferase assay in HEK293/ACE2/TMPRSS2 cells. As expected, the luciferase assay indicated that rXEC-infection activated NF-κB promoter more than rJN.1- and rXEC/N:P204R- infection in HEK293/ACE2/TMPRSS2 cells^44^ (**Figure 4D**).

To assess NF-κB-induced inflammatory signaling provoked by viral infection *in vivo*, we extracted mRNA of the lung hilum and periphery areas at 2 d.p.i., and quantified the expression of four interferon-stimulated genes (ISGs) (*Il-1β, Il-6, Il-8*, and *Ccl2*) (**Figure 4E**). In the lung hilum, rXEC-infection upregulated *Il-6 and Il-8* expression more than rJN.1- and rXEC/N:P204R-infection, suggesting that XEC could induce severe inflammation, supporting the histopathological data. In the lung hilum, the expression levels of *Il-1β* and *Ccl2* were also higher in rXEC- infected lungs than in rJN.1-infected lungs. Taken together, these results indicate that the N:R204P mutation enhances inflammation through NF-κB activation.

## Conclusion

In this study, we elucidated the properties of the Omicron XEC variant by investigating the growth kinetics *in vitro* and *in vivo*, resistance to antiviral drugs, fusogenicity of the S protein, and intrinsic pathogenicity. Epidemic dynamics modeling showed that two hallmark mutations in the XEC lineage, S:T22N and S:F59S, contribute significantly to its rapid spread. To date, Omicron subvariants had mainly altered the function of S protein to avoid immune pressure in the human population.^6–15^ However, recent studies suggest that the evolutionary potential of SARS-CoV-2 via S protein may be approaching a limit.^15,45–48^ For example, JN.1 subvariant demonstrates enhanced immune evasion due to S:L455S mutation, but this comes at the cost of reduced replication efficiency and diminished lung pathogenicity.^15,45^ Our data indicate that the acquisition of the N:R204P mutation, which impairs the function of N to inhibit NF-κB activation, enhances pathogenicity of XEC compared to JN.1. Thus, the strategies to modulate certain virological properties by mutations in non-spike proteins may become more common in the future evolution of SARS-CoV-2. In addition, close surveillance of XEC lineage is warranted, as it possesses the potential to become a dominant variant with both enhanced immune evasion and pathogenicity, driven by recombination.

## METHOD DETAILS

### Ethics statement

All experiments with hamsters were performed in accordance with the Science Council of Japan’s Guidelines for the Proper Conduct of Animal Experiments. The protocols were approved by the Institutional Animal Care and Use Committee of National University Corporation Hokkaido University (approval ID: 20-0123).

### Virus-like particle assay

A virus-like particle (VLP) assay was employed as a physiological model to test the efficiency of packaging and assembly as a function of the mutations on SARS-CoV-2 N protein.^32,33^ The assay was conducted as described previously.^32,33^ Briefly, the VLPs were generated by co-expressing all four structural proteins of SARS-CoV-2 in HEK293T cells along with a construct containing ∼1-kb viral packaging signal PS9 incorporated into the untranslated region of a firefly luciferase reporter. VLPs in the supernatant carrying luciferase reporter were added to receiver HEK293T cells stably expressing ACE2 and TMPRSS2 (HEK293T/ACE2/TMPRSS2 cells) in 96-well plates. Luminescence in receiver cells was measured at 12-16 hours post infection using luciferase assay system (Promega).

### Cell culture

HEK293 (ATCC) and HEK293T (ATCC) cells were maintained in DMEM (high glucose) containing 10% FBS and 1% PS. VeroE6/TMPRSS2 cells (VeroE6 cells stably expressing human TMPRSS2; JCRB Cell Bank)^27^ were maintained in DMEM (low glucose) containing 10% FBS, G418 (1 mg/ml), and 1% PS. Calu-3 cells (ATCC) were maintained in Eagle’s minimum essential medium (EMEM) containing 10% FBS and 1% PS. Calu-3/DSP1-7 cells (Calu-3 cells stably expressing DSP1-7^49^ were maintained in EMEM (Wako, Cat# 056-08385) containing 20% FBS and 1% PS.

### NF-κB promoter assay

HEK293/ACE2/TMPRSS2 cells were seeded onto 24-well plates and transfected. To assess NF-κB signaling, 50ng of the pNL3.2.NF-κB-RE vector (Promega) was utilized. Following transfection for 24 h, cells were infected for 1 h. The cells were then lysed using a passive lysis buffer and subjected to luciferase activity measurements by the Luciferase Reporter Assay System using an AB-2270 Luminescencer Octa (Atto).

### Modeling the relationship between amino acid mutations and epidemic dynamics

To identify the mutations contributing to the rapid spread of XEC, we utilized a hierarchal Bayesian multinomial logistic model previously developed by our research group to estimate the effects of mutations on fitness.^10^ Here, fitness refers to the relative Re between variants, estimated under the assumption that the relative values of the Re among variants remain constant over time. Unlike the conventional method of estimating Re using a multinomial logistic model,^26^ this model does not directly estimate the fitness of variants. Instead, it introduces a hierarchical structure to estimate fitness as a linear combination of mutations. As a result, this model can estimate not only the fitness of each variant but also the effects of individual mutations on fitness. For highly co-occurring mutations (mutation clusters), our method does not permit the estimation of individual effects, and thus, effects are estimated at the mutation cluster level. For details on the model, please refer to Ito et al.^10^ In this study, we estimated the effects of mutations not only in the spike protein but across all viral proteins.

The data used in this analysis were downloaded from the GISAID database (https://www.gisaid.org/) on January 26, 2025.^50^ For quality control, we excluded the data of viral sequences with the following features from the analysis: i) a lack of collection date information; ii) sampling in animals other than humans; iii) >1% undetermined nucleotide characters; or iv) sampling by quarantine. Furthermore, in this analysis, we analyzed viral sequences collected in the USA from January 1, 2024, to December 31, 2024.

For the analysis, we selected mutations (including substitutions, insertions, and deletions) observed in ≥100 sequences in the dataset we used. We then excluded mutations commonly (≥95%) detected in sequences analyzed.

According to the criteria above, 209 mutations were retrieved. Subsequently, we classified viral sequences into haplotypes, a group of viral sequences sharing the same set of mutations, according to the profile of the selected mutations.

We excluded haplotypes with ≤30 sequences from the downstream analyses. According to the criterion above, 366 types of haplotypes, composed of 51,127 sequences, were retrieved. Then, we clustered highly co-occurring mutations (i.e., a pair of mutations with >0.9 Pearson’s correlation in the mutation profile matrix) into mutation clusters. Consequently, our dataset included the profile of 209 mutation clusters for 366 haplotypes. For the reference haplotype, which is estimated to have a relative Re of 1, we selected the haplotype with the highest sequence count, corresponding to the major haplotype of JN.1.

Parameter estimation was performed via the MCMC approach implemented in CmdStan v2.30.1 (https://mc-stan.org) with CmdStanr v0.5.3 (https://mc-stan.org/cmdstanr/). Four independent MCMC chains were run with 500 and 2,000 steps in the warmup and sampling iterations, respectively. We confirmed that all estimated parameters showed <1.01 R-hat convergence diagnostic values and >200 effective sampling size values, indicating that the MCMC runs were successfully convergent. The above analyses were performed in R v4.2.1 (https://www.r-project.org/). Information on the estimated effect size of each mutation cluster on relative Re is summarized in **Table S1**.

### Plasmid construction

The nine pmW118 plasmids containing the partial gene of SARS-CoV-2 JN.1 were previously generated.^15,51^ To generate the recombinant XEC viruses, the mutations were introduced into the corresponding plasmids encoding JN.1 gene by inverse fusion PCR cloning. Sequences of all the plasmids used in this study were confirmed by a SeqStudio Genetic Analyzer (Thermo Fisher Scientific) and an outsourced service (Fasmac). Primer and plasmid information can be provided upon request.

### SARS-CoV-2 S-based fusion assay

A SARS-CoV-2 S-based fusion assay was performed as previously described.^24,52^ Briefly, on day 1, effector cells (i.e., S-expressing cells) and target cells (Calu-3/DSP1-7 cells) were prepared at a density of 0.6–0.8×10^6^ cells in a 6-well plate. On day 2, for the preparation of effector cells, HEK293 cells were cotransfected with the S expression plasmids and pDSP ^53^ using TransIT-LT1 (Takara). On day 3, 16,000 effector cells were detached and reseeded into a 96- well black plate (PerkinElmer), and target cells were reseeded at a density of 1,000,000 cells/2 ml/well in 6-well plates. On day 4, target cells were incubated with EnduRen live cell substrate (Promega) for 3 h and then detached, and 32,000 target cells were added to a 96-well plate with effector cells. Renilla luciferase activity was measured at the indicated time points using Centro XS3 LB960 (Berthhold Technologies). For measurement of the surface expression level of the S protein, effector cells were stained with rabbit anti-SARS-CoV-2 S S1/S2 polyclonal antibody (Thermo Fisher Scientific, 1:100). Normal rabbit IgG (Southern Biotech, 1:100) was used as a negative control, and APC-conjugated goat anti-rabbit IgG polyclonal antibody (Jackson ImmunoResearch, 1:50) was used as a secondary antibody. The surface expression level of S proteins was measured using CytoFLEX Flow Cytometer (Beckman Coulter) and the data were analyzed using FlowJo software v10.7.1 (BD Biosciences). For calculation of fusion activity, Renilla luciferase activity was normalized to the mean fluorescence intensity (MFI) of surface S proteins. The normalized value (i.e., Renilla luciferase activity per the surface S MFI) is shown as fusion activity.

### Antiviral drug assay using SARS-CoV-2 clinical isolates and human iPSC-derived lung organoids

The antiviral drug assay was performed as previously described. The human iPSC-derived lung organoids were infected with either JN.1 or XEC isolate (100 TCID50) at 37 °C for 2 h. Following infection, the cells were washed with DMEM and cultured in DMEM supplemented with 10% FCS, 1% PS, and the serially diluted EIDD-1931 (an active metabolite of Molnupiravir; Cell Signaling Technology), Nirmatrelvir (MedChemExpress), Remdesivir (Clinisciences), or Ensitrelvir (MedChemExpress). At 72 h post-infection, the culture supernatants were collected, and viral RNA was quantified using RT-qPCR. The assay of each compound was performed in triplicate, and the 50% effective concentration (EC50) was determined using Prism 9 software v9.1.1 (GraphPad Software).

### SARS-CoV-2 preparation and titration

The working virus stocks of SARS-CoV-2 were prepared and titrated as previously described. In this study, clinical isolates of JN.1 (strain LG0688; GISAID ID: EPI_ISL_18771637) and XEC (strain TKYnat18145; GISAID ID: EPI_ISL_19512397) were used.

Recombinant viruses were generated by a circular polymerase extension reaction (CPER).^44^ The resultant CPER products were transfected into VeroE6/TMPRSS2 cells as described previously.^12^ All the viruses were stored at −80°C until use and viral genome sequences were confirmed by SANGER sequencing as described above.

### Titration and growth kinetics

The infectious titers in culture supernatants obtained from the infected cells were determined by quantifying the 50% tissue culture infectious dose (TCID50).^54^ For growth kinetics, the viruses were respectively inoculated into VeroE6/TMPRSS2 cells in 12-well plates at a multiplicity of infection (MOI) of 0.01. The infectious titers of the indicated timepoints were determined.

### Airway-on-a-chip

Airways-on-a-chip were prepared as previously described.^55^ Human lung microvascular endothelial cells (HMVEC-L) were obtained from Lonza and cultured with EGM-2-MV medium (Lonza). For preparation of the airway-on-a-chip, the bottom channel of a polydimethylsiloxane (PDMS) device was first precoated with fibronectin (3 μg/ml, Sigma-Aldrich). The microfluidic device was generated according to our previous report.^56^ HMVEC-L cells were suspended at 5,000,000 cells/ml in EGM2-MV medium. Then, 10 μl of suspension medium was injected into the fibronectin-coated bottom channel of the PDMS device. The PDMS device was turned upside down and incubated. After 1 h, the device was turned over, and EGM2-MV medium was added into the bottom channel. After 4 d, airway organoids (AO) were dissociated and seeded into the top channel. Aos were generated according to our previous report.^57^ AOs were dissociated into single cells and then suspended at 5,000,000 cells/ml in the AO differentiation medium. Ten microliters of suspension medium were injected into the top channel. After 1 h, the AO differentiation medium was added to the top channel. In the infection experiments, the AO differentiation medium, containing either recombinant viruse (500 TCID50), was inoculated into the top channel. At 2 h.p.i., the top and bottom channels were washed and cultured with AO differentiation and EGM2-MV medium, respectively. The culture supernatants were collected, and viral RNA was quantified using RT-qPCR.

### Assessment of viral pathogenicity in hamsters

Animal experiments were performed as previously described.^6–15^ In brief, Syrian hamsters (male, 4 weeks old) were intranasally inoculated under anesthesia with the viruses (2,000 TCID50 in 100 μL for hamsters) or saline (100 μL). Body weight was recorded daily by 7 d.p.i. Lung tissues were anatomically collected at 2 and 5 d.p.i. The viral RNA load in the respiratory tissues was determined by RT-qPCR as described previously.^58^ These tissues were also used for IHC and histopathologic analyses as previously described.^6–15^ The viral proteins were visualized by anti-SARS-CoV-2 N monoclonal antibody (R&D Systems, 1:400). Pathological features—including (i) bronchitis or bronchiolitis, (ii) hemorrhage with congestive edema, (iii) alveolar damage with epithelial apoptosis and macrophage infiltration, (iv) hyperplasia of type II pneumocytes, and (v) the area of hyperplasia of large type II pneumocytes—were evaluated by certified pathologists after H&E staining. Images were incorporated as virtual slides by NDP.scan software v3.2.4 (Hamamatsu Photonics). The area of N-protein positivity and inflammation was measured using Fiji software v2.2.0 (ImageJ).

To evaluate inflammation levels evoked by viral infection in hamsters, 500 μg of the lung RNA was used to synthesize cDNA with SuperScript IV VILO Master Mix. The resulting cDNA was used to quantify the expression of host genes^59,60^ with a Power SYBER Green Master Mix (Thermo Fisher Scientific) and QuantStudio Real-time PCR System (Thermo Fisher Scientific).

### Quantification and statistical analysis

Statistical significance was tested by one-way ANOVA with Tukey’s multiple comparisons test using GraphPad Prism 9 unless otherwise noted. The values p < 0.05 were considered statistically significant (^∗^*p* < 0.05, ^∗∗^*p* < 0.01, ^∗∗∗^*p* < 0.001, ^∗∗∗∗^*p* < 0.0001). In the time-course experiments, a non-parametric permutation test was performed to evaluate the difference between experimental conditions thorough all timepoints. For each comparison, the area under the curve (AUC) was calculated as the sum of the values across timepoints. Group labels were randomly shuffled to generate the null distribution of AUC differences, and two-sided *P* values were calculated based on this distribution. Subsequently, familywise error rates (FWERs) were calculated by the Benjamini-Hochberg method. These analyses were performed in R v4.2.1 (https://www.r-project.org/). All assays were performed independently at least 2 times.

## CONSORTIA

The Genotype to Phenotype Japan (G2P-Japan) Consortium: H. Sawa, K. Matsuno, N. Nao, K. Shishido, H. Ito, Y. Kaku, N. Misawa, A. Plianchaisuk, Z. Guo, A. Hinay Jr., K. Uriu, Y. Kosugi, S. Fujita, J. M. Tolentino, L. Chen, L. Pan, M. Suganami, M. Chiba, K. Yasuda, K. Iida, N. Ohsumi, K. Yoshimura, K. Sadamasu, M. Nagashima, H. Asakura, I. Yoshida, S. Nakagawa, A. Takaori-Kondo, K. Shirakawa, K. Nagata, R. Nomura, Y. Horisawa, Y. Tashiro, Y. Kawai, R. Hashimoto, Y. Watanabe, A. Sakamoto, N. Yasuhara, T. Hashiguchi, T. Suzuki, K. Kimura, J. Sasaki, Y. Nakajima, H. Yajima, T. Irie, R. Kawabata, K. Tabata, B. MST Monira, R. Shimizu, M. Jonathan, Y. Mugita, O. Takahashi, T. Ueno, M. Toyoda, A. Saito, M. Shofa, Y. Shibatani, and T. Nishiuchi.

## ACKNOWLEDGMENTS

We would like to thank all members of The Genotype to Phenotype Japan (G2P- Japan) Consortium. We thank H. Kubo, M. Tetsuka, S. Shimamura, K. Yano, and A. Hisanaga for their secretory work and H. Murota for his technical assistance. We also thank Tokyo Metropolitan Institute of Public Health (Tokyo, Japan) for providing a clinical isolate. We gratefully acknowledge the numerous laboratories worldwide that have provided sequence data and metadata to GISAID.

This study was supported in part by AMED SCARDA Japan Initiative for World-leading Vaccine Research and Development Centers “UTOPIA” (JP223fa627001, to K.S.), AMED SCARDA Program on R&D of new generation vaccine including new modality application (JP223fa727002, to K.S.); AMED SCARDA Hokkaido University Institute for Vaccine Research and Development (HU-IVReD) (223fa627005h0001, to T.F.); AMED Project for Advanced Drug Discovery and Development (JP21nf0101627, to T.F.); AMED Research Program on Emerging and Re-emerging Infectious Diseases (JP21fk0108493, to T.F.; JP22fk0108617, to T.F.; JP22fk0108516, to T.F.; JP22fk0108146, to K.S.; JP21fk0108494 to G2P-Japan Consortium, Shinya T., T.F., and K.S.); AMED Research Program on HIV/AIDS (JP22fk0410039, to K.S.; JP22fk0410055, to T.I.); AMED CREST (JP21gm1610005, to K.T.; JP22gm1610008, to T.F.); JST PRESTO (JPMJPR22R1, to J.I.); JST CREST (JPMJCR20H4, to K.S.); JSPS KAKENHI Fund for the Promotion of Joint International Research (International Leading Research) (JP23K20041 to K.S., and T.F.); JSPS KAKENHI Grant-in-Aid for Scientific Research B (21H02736, to T.F.); JSPS KAKENHI Grant-in-Aid for Scientific Research C (22K07103, to T.I.); JSPS KAKENHI Grant-in-Aid for Early-Career Scientists (20K15767, to J.I.; 22K16375, to H.N.); JSPS Core-to-Core Program (A. Advanced Research Networks) (JPJSCCA20190008, to K.S.); The Cooperative Research Program (Joint Usage/Research Center program) of Institute for Life and Medical Sciences, Kyoto University (to K.S.); the Joint Research Program of Institute for Genetic Medicine, Hokkaido University (to K.Y., and T.T.); International Joint Research Project of the Institute of Medical Science, the University of Tokyo (to T.I.); Akiyama Life Science Foundation (to T.T.); Japan Antibiotics Research Association (to T.T.); Hirose Foundation (to T.T.); The Tokyo Biochemical Research Foundation (to K.S.); Takeda Science Foundation (to R.S., T.F., and T.I.); The Uehara Memorial Foundation (to T.I.); Tobe Maki Foundation (to S.S.); Hokkaido University Support Program for Frontier Research (to T.F.); and Tsuchiya Mitsubishi Foundation (to K.S.). M.O. gratefully acknowledges support for this study from NIH U19 AI135990, Roddenberry Foundation, P. and E. Taft, James B. Pendleton Charitable Trust. M.O. is a Chan Zuckerberg Biohub – San Francisco Investigator.

## AUTHOR CONTRIBUTIONS

Shuhei T. and S.D. performed cell culture experiments. Shuhei T., T.T., and K.Y. performed animal experiments. M.T., L.W., Y.O., and Shinya T. performed histopathological analysis. S.D., and K.T. prepared human lung organoids and airway-on-a-chip systems. S.D., and K.T. performed antiviral drug tests. H.N., and T.I. performed SARS-CoV-2 S-mediated membrane fusion assays. T.T. and M.O. designed, performed, and interpreted the VLP experiments. Shuhei T. generated recombinant viruses. J.I. performed phylogenetic and bioinformatics analyses. Shuhei T. performed statistical analyses. Shuhei T., K.S., Shinya T., T.T., and T.F. designed the experiments and interpreted the results. Shuhei T., T.T., and T.F. wrote the original manuscript. All authors reviewed and proofread the manuscript.

## Supplementary files

Supplementary Table 1. Estimated effect of each substitution on Re estimated by a hierarchal Bayesian multinomial logistic model

**Supplementary Figure 1.**
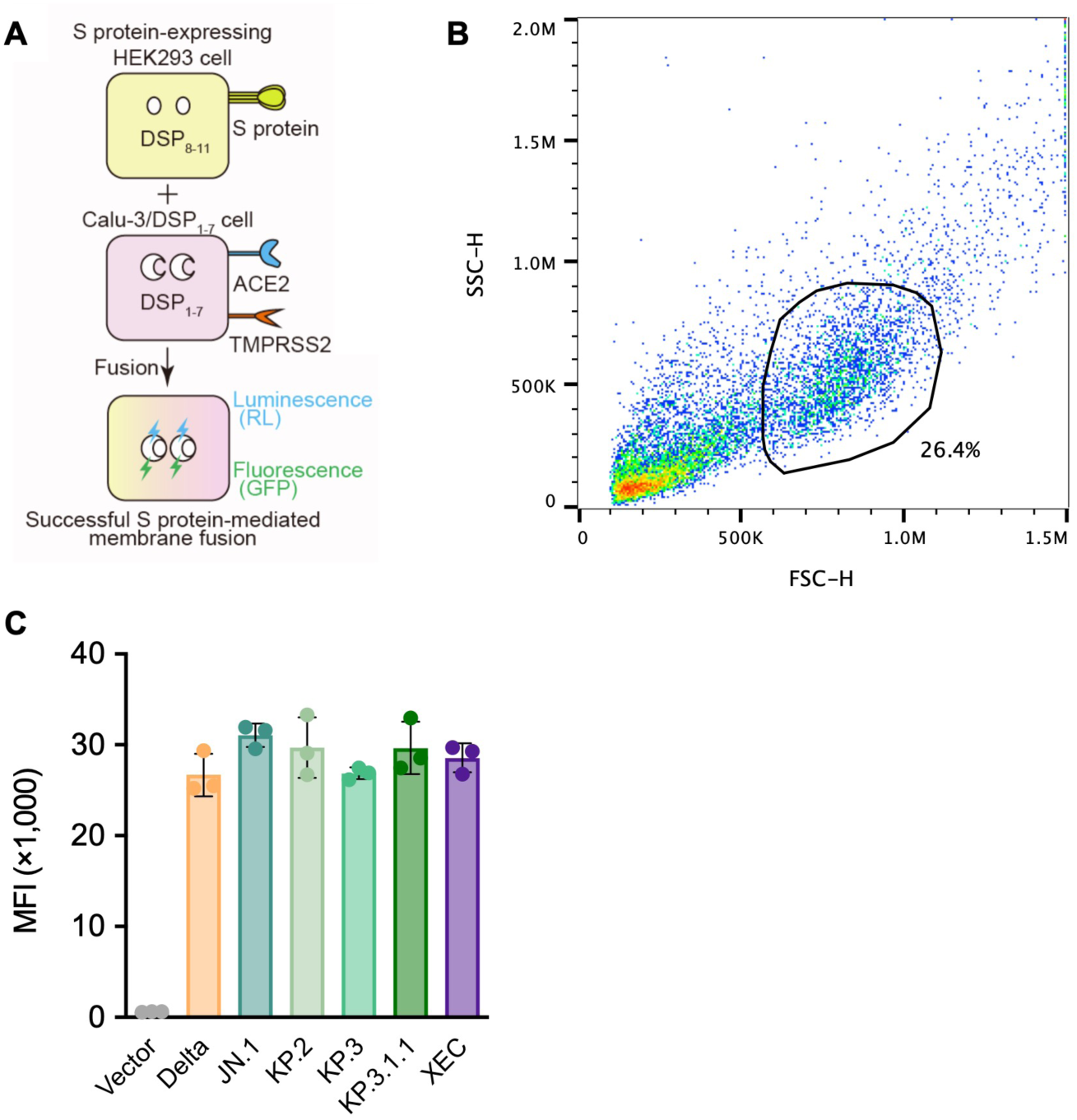
Virological features of XEC S, related to Figure 2A (A) Schematic diagram of SARS-CoV-2 S-mediated membrane fusion assay using Calu-3/DSP cells. (B) Gating strategy for flow cytometry of S protein expressing cells. (C) S protein expression on the cell surface. Mean fluorescence intensity (MFI) of surface S protein by flow cytometry. The summarized data are shown.

**Supplementary Figure 2.**
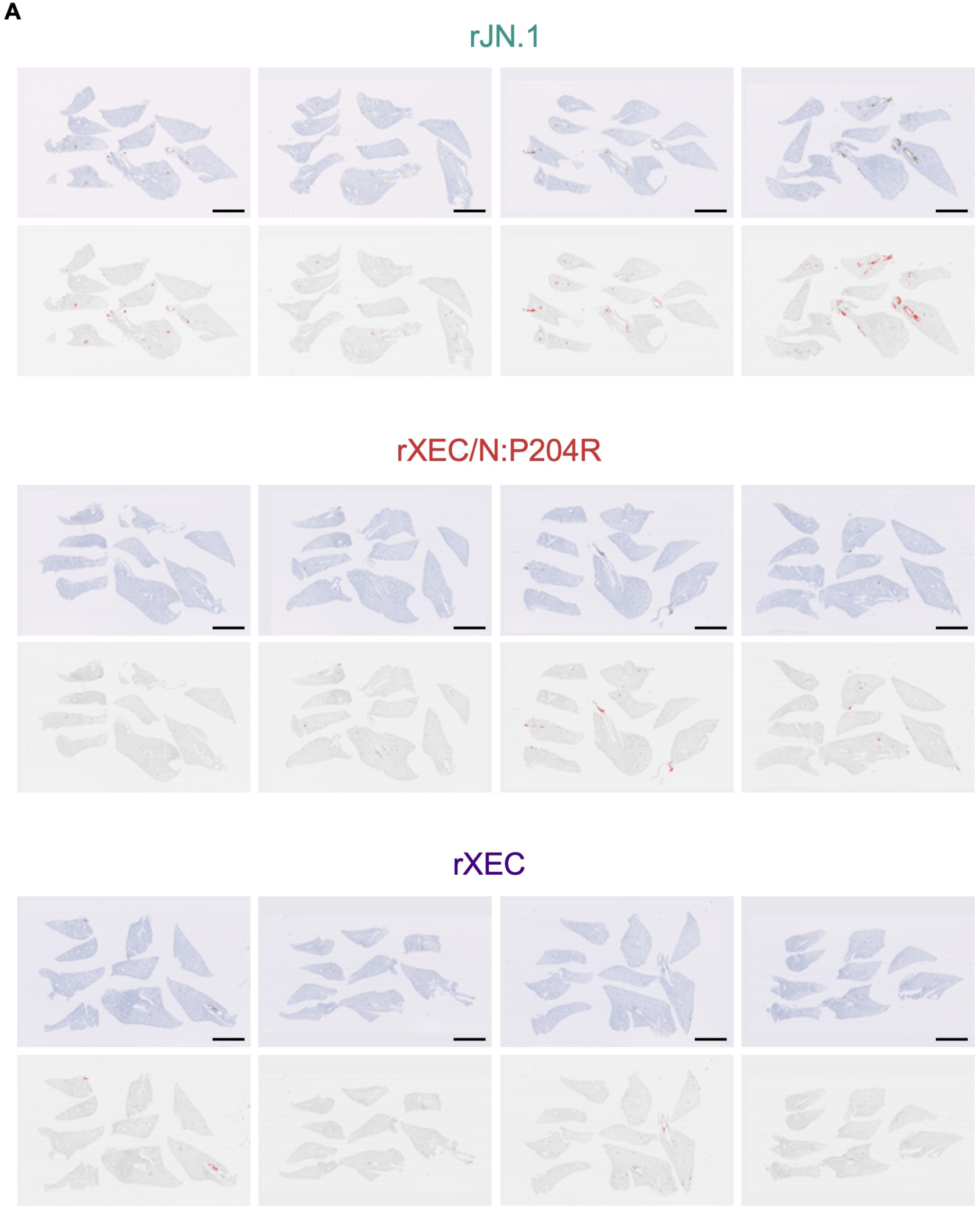
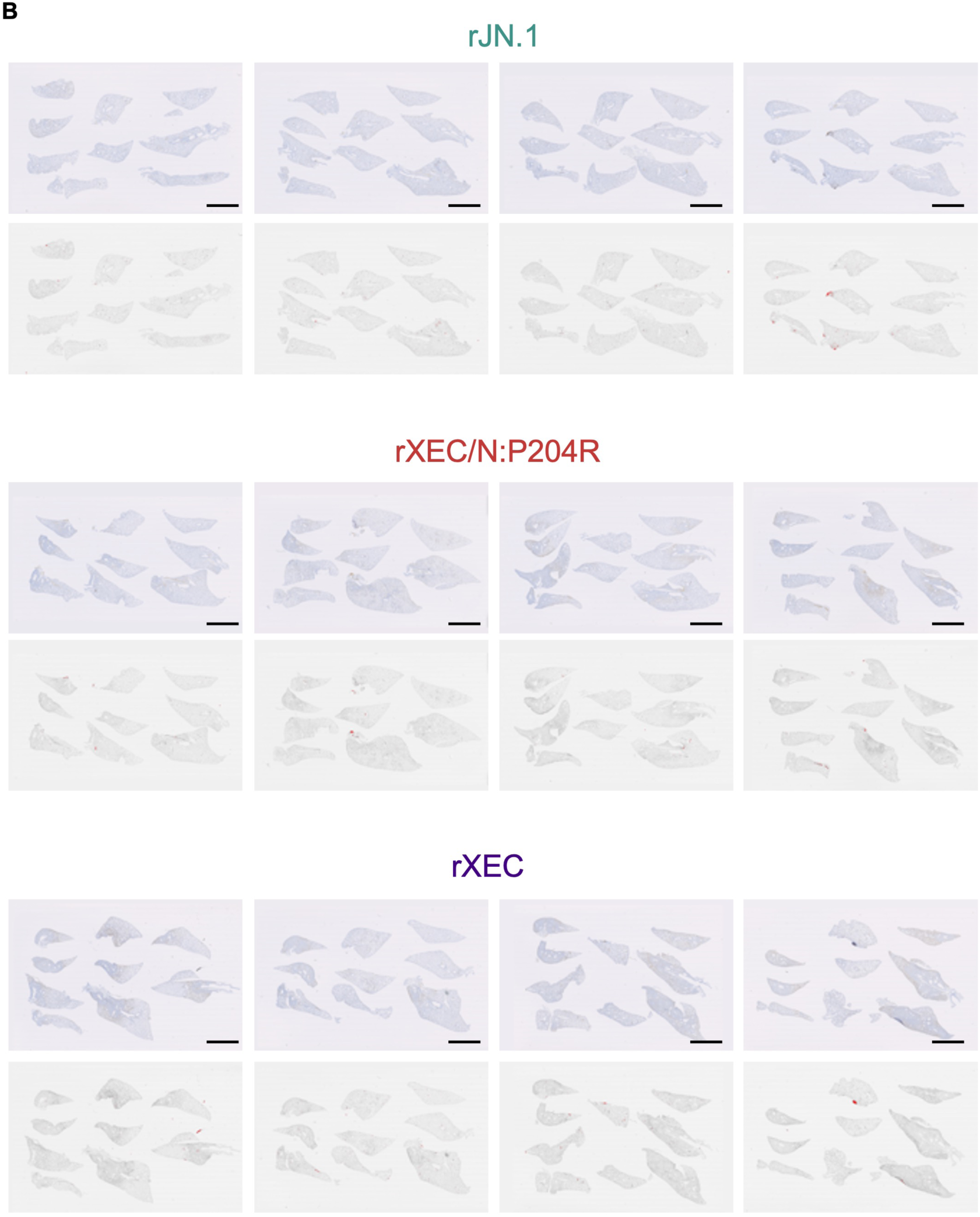
Histological observations in infected hamsters, related to Figure 4A (A and B) Immunohistochemistry of the viral N protein in the lungs of infected hamsters at 2 d.p.i (A) and 5 d.p.i (B) The percentages of N-positive cells in whole lung lobes are shown in the lower panels for each infection. Scale bars, 5 mm.

**Supplementary Figure 3.**
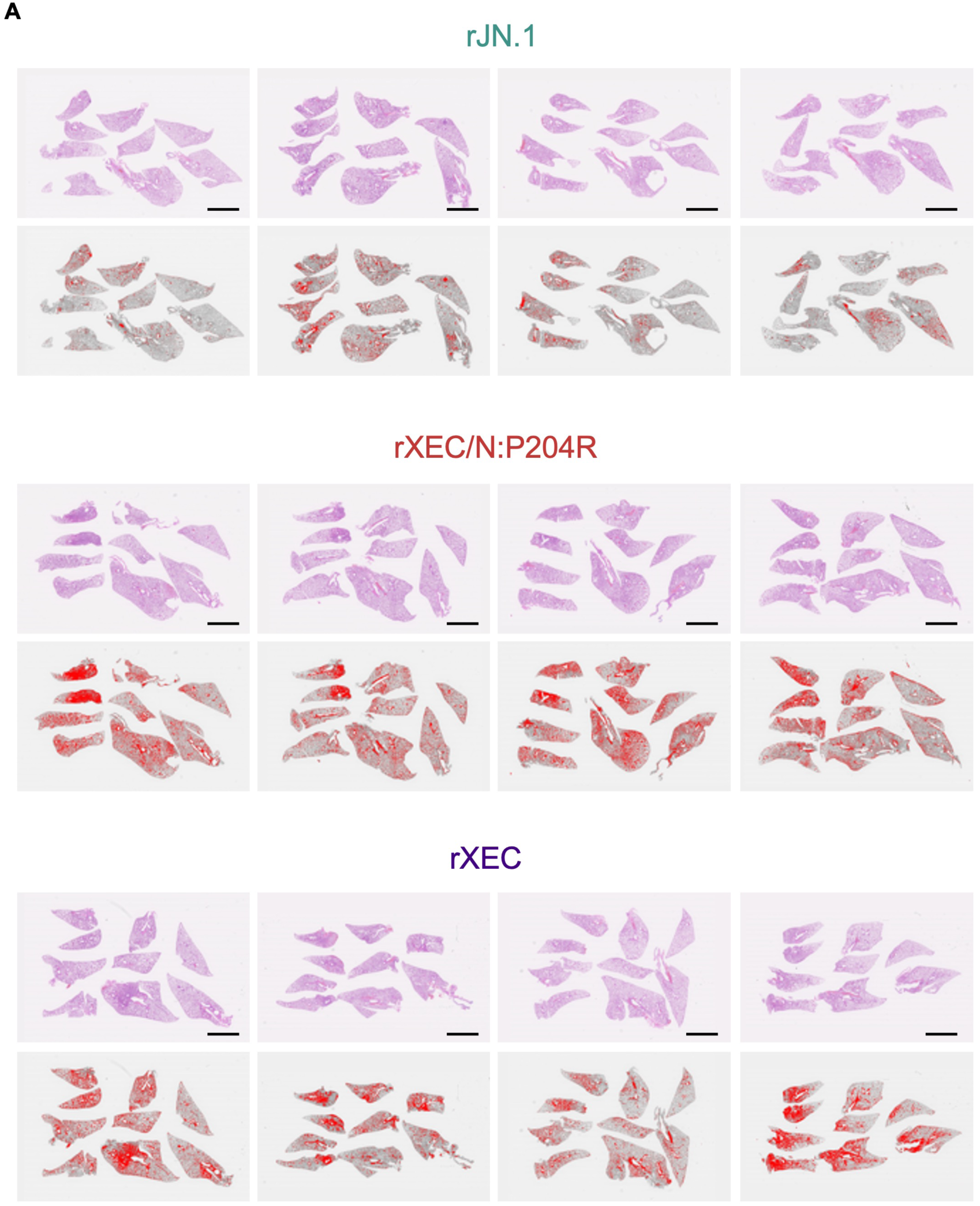
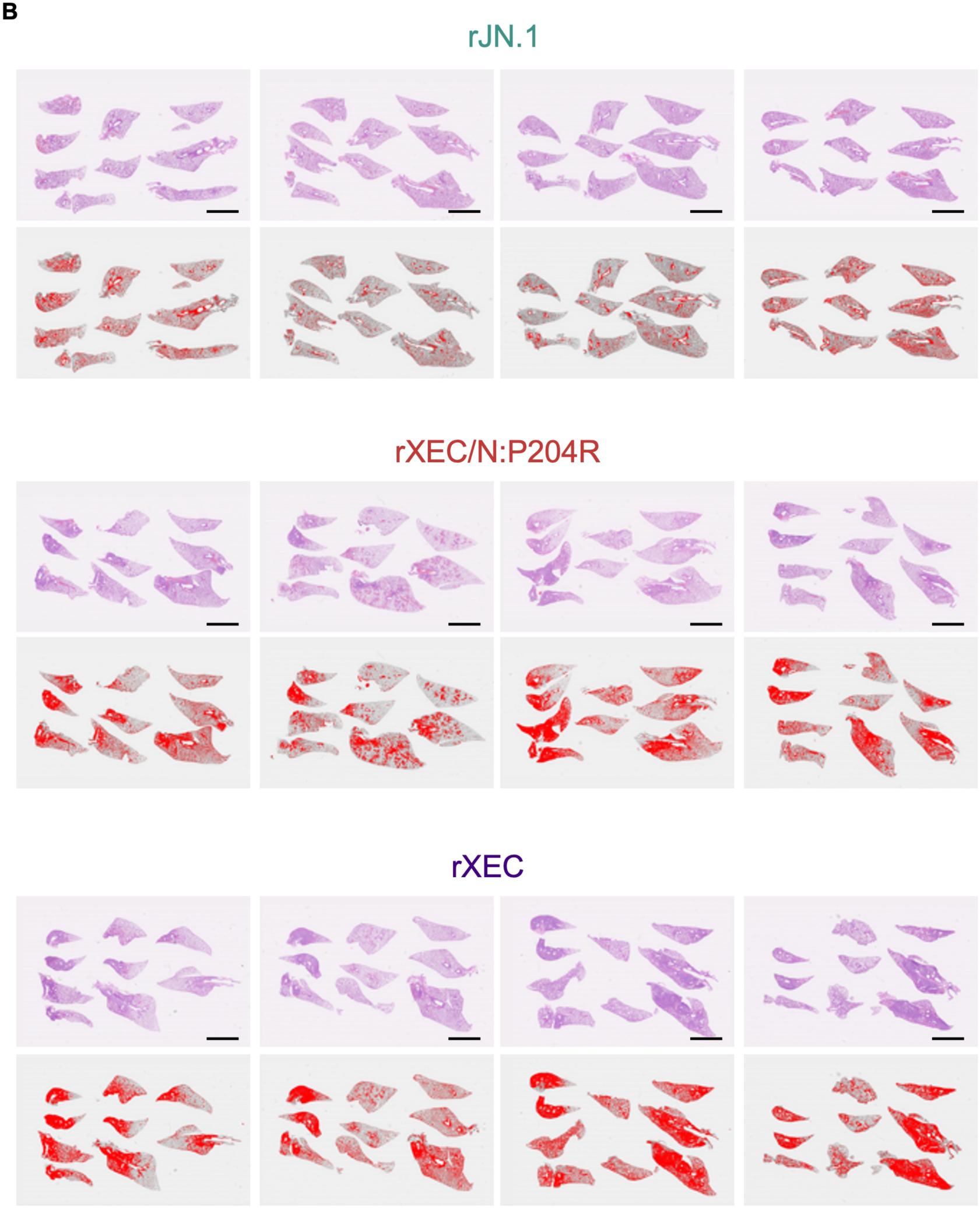
Histological observations in infected hamsters, related to Figure 4B (A and B) Hematoxylin and eosin staining of the lungs of infected hamsters at 2 d.p.i. (A) and 5 d.p.i. (B) Inflammatory areas with type II pneumocytes are shown in red and the percentage calculated in the lower panels for each infection. Scale bars, 5 mm.

